# ASSESSING POST-FIRE VEGETATION TRAJECTORIES USING MACHINE LEARNING AND REMOTE SENSING: EVIDENCE FROM A MEDITERRANEAN SITE

**DOI:** 10.64898/2026.05.26.727844

**Authors:** Emiliano Agrillo, Nazario Tartaglione, Alessandro Mercatini, Alice Pezzarossa, Gianluigi Ottaviani, Mara Baudena, Federico Filipponi

**Affiliations:** Italian National Institute for Environmental Protection and Research (ISPRA), Roma, Italy; Research Institute on Terrestrial Ecosystems, National Research Council (CNR-IRET), Porano, Italy - National Biodiversity Future Center (NBFC), Palermo, Italy; Climate, National Research Council (CNR-ISAC), Turin, Italy - National Biodiversity Future Center (NBFC), Palermo, Italy; Institute of Environmental Geology and Geoengineering, National Research Council (CNR-IGAG), Montelibretti, Italy

**Keywords:** Fire disturbance, Landscape recovery, Mediterranean ecosystems, Post-fire vegetation trajectories, Spatial-Temporal monitoring

## Abstract

Fire has acted as a major eco-evolutionary force since the evolutionary appearance of plants, shaping plant-traits, diversity, vegetation assembly, and ecosystem functioning. Its ecological role depends on long-term fire regimes. Anthropogenic land-use change and climate warming are disrupting these regimes, particularly in densely populated regions such as the Mediterranean Basin. In the Italian peninsula (Mediterranean region) fire activity peaks during the dry summer months and is projected to intensify under climate change scenarios.

Recent methodological developments – based on emerging satellite data, ground-based observations combined with Random Forest (RF) habitat classification, and spectral indices such as the NDVI provide a robust framework for monitoring post-fire land-cover dynamics over time.

In this study, we applied RF modelling to classify vegetation cover using a 2017–2024 satellite imagery time series of the Monte Pisano area (central Italy) to assess pre- and post-fire vegetation trajectories. Evergreen shrubs and trees exhibited rapid post-fire regrowth, whereas coniferous stands showed slower recovery rates. NDVI trends revealed an expected sharp decline immediately after the fire, followed by gradual recovery of broadleaf forests and shrubland communities. Moreover, our results indicated a progressive increase in the cover of native deciduous and evergreen species of high conservation value (listed under the Habitat Directive).

The framework delivers spatially explicit insights into post-fire recovery, supporting targeted management, restoration under European Nature Restoration Regulation, and long-term monitoring in Mediterranean ecosystems. Incorporating fine-scale environmental variables may further improve classification accuracy and enhance assessments of vegetation resilience and ecosystem recovery following fire events.

**Highlights:**

- Recurring fires strongly affect ecosystem structure and function in Mediterranean landscapes.
- Integrating remote sensing with Random Forest models enables effective monitoring of post-fire vegetation recovery over time.
- NDVI time series provide reliable proxies for tracking vegetation vigor and land-cover change.
- Post-fire recovery trajectories are shaped by fire severity, vegetation physiognomy, plant functional types, and soil conditions.
- Targeted restoration and management interventions informed by spatial–temporal vegetation patterns are urgently needed.
- The proposed framework aligns with objectives of the EU Nature Restoration Regulation for ecosystem and habitat recovery.

## 1 Introduction

Fire has been an eco-evolutionary force since the appearance of terrestrial plants on Earth (Bowman et al., 2009; Pausas & Keeley, 2014; Pausas et al., 2018; Pausas et al., 2025), because plant material constitutes the fuel for any wildland fire to occur. As such, fire has largely contributed to shape plant species’ forms, functions, diversity, distribution and assembly of vegetation types, with direct and indirect impacts on ecosystem functioning (e.g., carbon and nutrient cycling). In this eco-evolutionary context, the key factor is not whether or not fire occurs in a given landscape or region (some are more or less prone to this disturbance agent, e.g., Lamont & He, 2017; Pausas & Keeley, 2017) rather its long-term regime, defined by the predictable frequency, intensity, extent, seasonality and type (Archibald et al., 2013). In this context, recent anthropogenic alterations of land use and climate are generating major disruptions to these long-term dynamics, with potentially detrimental effects on various aspects of socio-ecological systems (Kelly et al., 2020; Lecina-Diaz et al., 2021). This is especially so in areas where human density, activities and pressures are high, as in Mediterranean basin, which constitutes a major concern for land managers dealing with ecosystem resilience and post-fire vegetation dynamics. Fire severity and intensity can strongly influence post-fire vegetation recovery, particularly in combination with other ecological factors (Viana-Soto et al., 2017). These include vegetation type (Keeley et al., 2011; Nolan et al., 2021), competitive native seed and bunk banks for the restoration of local-regional species pool (e.g., Ladouceur et al., 2018) and geomorphological conditions (Ice et al., 2004; McGuire et al., 2024).

Recent research highlights the effectiveness of remote sensing combined with machine learning techniques for assessing post-fire land-cover (LC) recovery (Priya & Vani, 2024). Algorithms – such as Random Forest (RF) – have been widely applied to classify vegetation cover of different habitat types (Agrillo et al., 2025) and their changes from high-resolution satellite imagery, enabling detailed, pixel-level mapping of postfire recovery (Kurbanov et al., 2022). Several studies have successfully employed RF with multitemporal imagery to characterize recovery dynamics across diverse ecosystems (Chen et al., 2011). The Normalized Difference Vegetation Index (NDVI) and its derivative, the delta-NDVI (ΔNDVI), are among the most widely used indices for capturing specific aspects of post-fire disturbance, as they provide robust proxies for photosynthetic activity of leaf biomass (Tucker, 1979; Pettorelli et al., 2005). Moreover, remote sensing mapping provides pivotal spatial information for assessing LC across regions and time periods (Wulder et al., 2003; Wulder et al., 2018). These datasets are widely used to monitor vegetation dynamics, evaluate ecosystem extent, and detect land degradation and disturbances (Mücher et al., 2009; Nagendra, 2013; Roelofsen et al., 2014). Land-cover trajectories following disturbance events, such as post-fire vegetation recovery, occur across multiple spatial and temporal scales (Foley et al., 2005; Wulder et al., 2020; Zhang et al., 2024). Effective monitoring of these changes requires approaches that can capture natural “patch dynamics” and trajectories over long time periods and across diverse spatial extents (Pickett, 1985; Wulder et al., 2018). Temporal analyses of post-fire vegetation recovery, stratified by ecosystem type and burn severity, can provide actionable insights for land managers, such as supporting identification and prioritization of areas for conservation or the development of effective post-fire management strategies.

Continuous LC change detection methods pose unique challenges. Many implementations rely on a single classifier trained at the beginning of the monitoring period, assuming that LC class characteristics remain stable over time (Demarquez et al., 2024). This assumption may be contentious in ecology, as spectral signatures can shift in response to successional dynamics, species turnover, or climate-driven phenological changes. Alternatively, periodically re-training classifiers (e.g., annually or seasonally) can account for temporal variability, but it increases computational demands and may reduce temporal consistency across maps (Zhang et al., 2022). Selecting between these strategies thus involves trade-offs between efficiency, accuracy, and ecological realism, highlighting the importance of tailoring LC change-detection frameworks to the dynamics of the system under study (Cheng et al., 2024).

Therefore, the primary strategies commonly applied in machine learning-based LC classification with satellite time-series data (Gómez, 2016) can be summarized as follows: (i) repeated model retraining to capture intra- and/or interannual variability (Franklin, 2015); and (ii) training a single model to generate a reference map that is subsequently projected using multitemporal satellite imagery (Wegman et al, 2016). This second approach can also be used to “backdate” maps, producing multi-year classifications from a single reference year using only updated spectral information (Pouliot, 2014). Both approaches have successfully employed the RF algorithm to assess vegetation trends over decadal periods. However, transferring models across time may introduce biases due to incomplete knowledge of local vegetation dynamics (Filippelli, 2024).

A key methodological gap remains to systematically comparing retrained and transferable models under changing environmental conditions. Addressing this gap is critical for robust LC monitoring that ensures both accuracy and long-term consistency. For these reasons, this study aims at evaluating the performance of two remote sensing–based modeling approaches for land cover classification and analysis of vegetation trajectories, which were tested over an eight-year period in a Mediterranean case-study site in central Italy, in an area unaffected by wildfire, before applying the best approach in a, wildfire-affected area. The choice of focusing on a restricted study area reflects research priorities that emphasize temporal rather than spatial transferability of the modeling approaches (Ludwig, 2022) and enables a rigorous evaluation of model performance through a structured validation framework.

We specifically set out to:

- Compare Random Forest classification performance using multi-year versus single-year training models;
- Assess post-fire land-cover trajectories;
- Quantify vegetation recovery patterns to guide targeted strategies for post-fire mediterranean landscape resilience and restoration.

## 2 Data and Methods

We used Random Forest models to predict post-fire land-cover trajectories. Two calibration strategies were first evaluated in an undisturbed site with comparable vegetation covers to identify the most accurate and stable approach. Models used environmental variables and Sentinel-2 metrics (2017–2024) to classify vegetation-cover types and assess post-fire recovery relative to unburned reference areas. Remotely sensed indices were used to assess vegetation vigor and characterize post-fire recovery dynamics in burned areas relative to unburned reference sites.

### 2.1 Study Area

The analysis focuses on natural and semi-natural Mediterranean ecosystems with documented fire histories (i.e from 1984 at present). For this reason, the Monte Pisano massif (917 m a.s.l.), an isolated subrange of the Apennines in central Italy (Tuscany), near the Tyrrhenian coast, was selected to classify LC and evaluate post-fire vegetation dynamics. The site was selected due to its natural vegetation post-fire dynamics, which has seen the area recolonized mainly by pre-existing native species (Bertacchi, 2022).

The landscape of Monte Pisano has been historically shaped by activities such as agricultural terracing, coppice felling, and seasonal grazing, which have influenced its current vegetation composition. The dominant vegetation cover consists primarily of maritime pine (*Pinus pinaster*) and holm oak (*Quercus ilex*) woodlands on the western slopes, while coppice chestnut (*Castanea sativa*) woodlands prevail in the northern and eastern regions (Bertacchi, 2023). Scattered patches of garrigue and evergreen shrubs occur throughout the area in various growth stages, likely in equilibrium with a regime of repeated fires (i.e., “pyroclimax” communities). Extensive olive groves occupy the transition zones between rural and natural areas, representing a traditional form of cultivation typical of the Mediterranean region. When abandoned, these groves can assume the characteristics of semi-natural ecosystems (Gennai-Schott, 2020) recolonized by the neighbouring phytocoenoses (e.g., evergreen plant communities).

Historically, fire has been the dominant disturbance agent in Monte Pisano, particularly affecting the western and southern slopes of massif over the past 50 years. Between 1970 and 1999, 59 fire events were recorded, with burned areas ranging from 5 to 600 ha (data from the Tuscany regional geo-portal GEOscopio). From 2000 to 2019, 16 additional fires occurred; the largest of these was at the end of September 2018 t in the Calci locality, which burned more than 1,000 ha.

Figure 1 shows the entire Monte Pisano study area, subdivided into burned (2000–2019) and unburned areas. The largest fire, which occurred in 2018 nearby the village of Calci, is also highlighted and constitutes the focus of the post-fire land-cover change analysis of this study.

**Figure 1.**
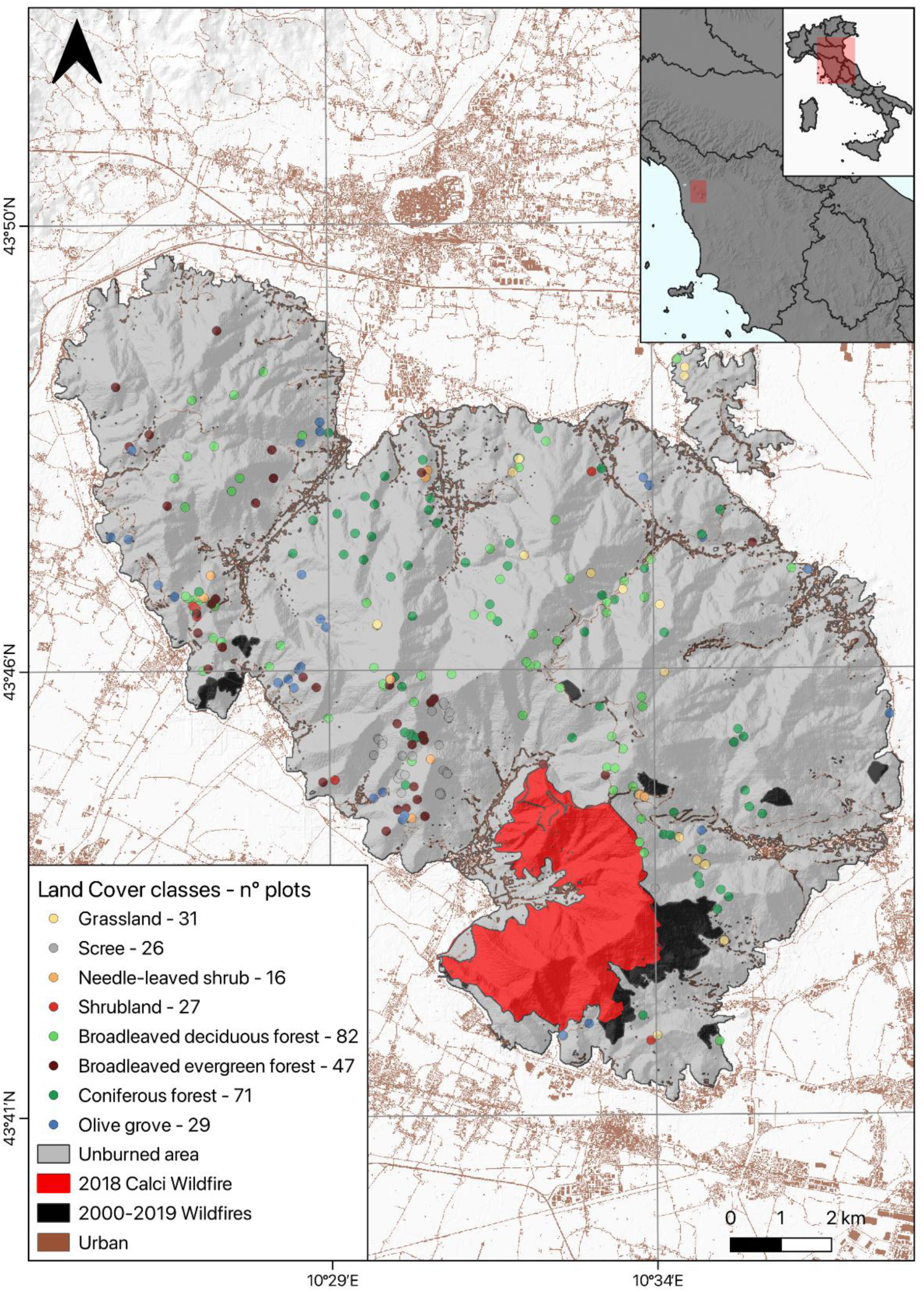
Study area map showing the 2018 Calci fire area (red), fires from 2000 to 2019 (black), and unburned area before the 2000 (grey). Dots indicate ground-truth observations for land-cover classes.

Therefore, across the entire study area, sites affected by fires after 2000 (Fig. 1) were excluded from the classification model training and are hereafter referred to as the “undisturbed reference area”. We assume that environmental drivers (i.e. geomorphological and climatic) within the reference area remained stable over the eight-year period considered in this work (2017–2024). Vegetation patterns in this portion of Monte Pisano are expected to be primarily shaped by natural dynamics (Trabaud, 1994; Allen, 2009). Other local disturbances, such as coppicing and pest outbreaks, were considered negligible, as their presence is locally minor and unlikely to affect the aims of the study.

### 2.2 Response and Predictor Variables for classification model

#### 2.2.1 Response Variables: target land-cover classes

The target (response) variables were obtained both from published georeferenced sampling datasets and field surveys conducted between 2023 and 2024, in order to increase the number of LC response observations within the study area. The number of target points was determined based on the extent of vegetation patterns within the region of interest, ensuring balanced data for RF model training and validation steps. A total of 329 ground-truth plots (60 from published sources and 269 from field observations) were collected within the Monte Pisano reference area (grey area in Figure 1), representing the main vegetation types identified by physiognomy and by occurrence of constant species (*sensu* Chytrý & Tichý 2003). The legend in Figure 1 shows the number of target points (i.e., plots) for the eight focal land-cover classes: grassland (G), scree (SC), needle-leaved shrub (N), shrubland (S), broadleaved deciduous forest (T1), broadleaved evergreen forest (T2), coniferous forest (T3), and olive groves (OG). The legend is based on EUNIS habitat definition (Chytrý et al., 2020), modified to reflect local land-cover classes (G, N, and OG).

Furthermore, to mitigate misclassification in the vegetation cover classification result arising from spatial autocorrelation, we implemented a stratified sampling strategy to ensure homogeneous and discrete vegetation plots (Legendre, 2012). Sampling was stratified by altitudinal belts and slope exposure, providing a robust framework for assessing model accuracy, reducing potential biases, and enabling a comprehensive evaluation of model performance (Li et al., 2021).

#### 2.2.2 Predictors: environmental and spectral data

A total of 182 variables were obtained through data mining and subsequently grouped into two main categories:

- <u>Environmental data:</u> 14 variables describing geomorphological, climatic, and soil characteristics. The geomorphological variables were selected for their relevance to plant dispersal (Immitzer 2019; Descombes 2020) and assumed to be time-invariant. Climatic data included mean precipitation and temperature. Soil properties (pH, depth to bedrock) were extracted from SoilGrids - 250 m resolution (Poggio et al. 2021);
- <u>Spectral data</u>: 168 reflectance values and indices derived from S2 Multi-Spectral Instrument. Spectral predictors consisted of S2 surface reflectance bands (distributed by Theia) and derived indices. Images from 2017–2024 with <90% cloud cover were used; monthly composites were generated by excluding pixels flagged as cloud, cloud shadow, snow, topographic shadow). Missing monthly values were imputed with the ‘*imputeTS’* R package (Moritz 2017). In addition to reflectance bands, four indices were calculated (i.e. EVI, NDYI, RI, CRI1), to capture pigment-related changes in vegetation during the vegetative and blooming peak as well as during senescence periods (Filipponi, 2022).

The final set of predictors consisted of annual time series derived from monthly S2 spectral bands and vegetation indices, integrated with a suite of environmental covariates - see Supporting Material (SM.1-Tables S1 & S2) for a detailed description of consulted sources, formulas used to compile and calculate spectral and environmental predictors, structured according to the objectives of this study.

### 2.3 LC Model Strategy and Stability Assessment

#### 2.3.1 Model Approaches

To ensure classifier reliability prior to application in fire-disturbed areas, calibration and stability assessment were conducted exclusively in an undisturbed reference site. This allowed evaluation of the temporal consistency of the RF model in a landscape with little or no short-term vegetation change. Two modeling approaches were compared in the Monte Pisano area, restricting analyses to areas unaffected by fires since 2000. Stable classifications under these conditions increase confidence in model performance in disturbed landscapes.

Therefore, the following land-cover classification strategies were implemented:

- <u>MULTI approach:</u> a separate RF model was trained each year using the S2 satellite imagery time-series for that specific year, combined with environmental predictors and response variables surveyed in 2023-2024. Each model was then applied to classify every pixel across the study area, producing LC maps from 2017 to 2024;
- <u>SINGLE approach</u>: a single RF model was trained using response variables, spectral, and environmental predictors for the reference year 2023. This model was then applied to predict LC spatial classifications for the years 2017–2024 without retraining; only the spectral data derived from S2 imagery were updated each year.

Both classification approaches were performed in *ranger* R package (Wright, 2017), an efficient implementation of the Random Forest algorithm (Breiman, 2001) particularly suited for high-dimensional data. The algorithm supports classification, regression, and survival analysis and includes features such as randomized trees (Geurts, 2006) and quantile regression forests (Meinhausen, 2006). All available predictor variables were retained as inputs, since RF is robust to multicollinearity among predictors (Dormann, 2013).

During the calibration stage, 80% of targets plots were used as training datasets and 20% as testing dataset for evaluating the prediction stage. The latter dataset serves as the actual label in cross-tabulation with predictions for a sample of occurrences from the mapped area. This process provides a definitive evaluation of both correct and incorrect classifications. The methods used for accuracy evaluation and area estimation depend on this cross-tabulation, which is referred to as an error or confusion matrix (Olofsson et al., 2014).

Overall accuracy or the percentage of accurately spatial prediction is the key map accuracy metric (Olofsson et al., 2014; Stehman and Foody, 2019). This single statistic measures map quality across all categories. Following the approach proposed by Olofsson et al. (2014), we calculated the standard error to assess accuracy and estimate consistency across random sample choices. Overall correctness can calculate user and producer accuracy (Story, 1986). The study generated user’s and producer’s accuracy estimates across years and land-cover classes, along with all overall annual accuracy values, each associated with a standard error (Olofsson et al., 2013).

#### 2.3.2 Model Stability Metrics

Using the outputs of the two applied approaches, we computed specific performance metrics to determine which approach more accurately represents vegetation dynamics within the study area and utilized those results to study post-fire LC trajectories.

We evaluated the spatial and temporal stability of classifications in areas unaffected by fire over the past 20–25 years (Trabaud, 1994). Therefore, fire-disturbed areas after 2000 were excluded. Therefore, to evaluate the most suitable approach for land-cover classification in both fire-affected and unaffected areas, we assessed the temporal stability of the model outputs using the following metrics:

- The coefficient of variation (*CV*) (Brown,1998) for each LC class. This metric enables a straightforward comparison of how variability differs across different classes over time. The *CV* is defined as:

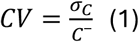 Where 𝜎_𝐶_ and 𝐶^-^ are, respectively, the standard deviation and mean computed over the years of the number of pixels of class *C*. The coefficient of variation (*CV*) provides a normalized measure of prediction consistency over time, enabling assessment of the temporal stability of predicted values. Lower CV values indicate more stable and reliable model predictions;
- The pixel stability measures (Boston et al, 2023) evaluating the number of changes of single pixels, normalized by the number of years. The parameter for pixel stability is calculated using the following equation:

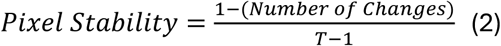 Where *T-1* represents the maximum number of changes. This gives a value between 0 and 1, where a value of 1 indicates no temporal change, whereas a value of 0 indicates the maximum temporal change.

### 2.4 Post-fire LC validation using ground-truth

We carried out a field survey in December 2024 to assess the accuracy of the 2023 model outputs within the 2018 wildfire-affected area in Calci. A total of 101 plots were selected using a stratified random sampling design, balanced across the predicted LC classes. Model performance was evaluated by comparing predicted and observed classes, with predictions classified as correct or incorrect based on membership in the corresponding LC class. Additionally, we generated a contingency table and computed the F1 score, defined as the harmonic mean of precision and recall (Sofaer et al., 2019), as well as the area under the curve (AUC) for multiclass prediction (Hand & Till, 2001). Specifically, we employed the ‘*multiclass.roc’* function from the R package ‘*pROC’* (Robin et al., 2011), which computes the average AUC by performing pairwise comparisons between classes.

### 2.5 Metrics to assess post-fire LC trajectories

Post-fire vegetation recovery in the 2018 burn area (1150 hectares) was assessed using satellite data (i.e. vegetation index). S2 multispectral imagery was processed to compute the delta-Normalized Difference Vegetation Index (ΔNDVI), a widely used metric for evaluating post-disturbance ecosystem trajectories (Gouveia et al., 2010; Lasaponara et al., 2022).

NDVI was calculated using the standard formulation:

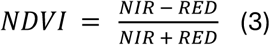

Where NIR and RED correspond to S2 bands, B08 (resolution: 10 m/px; central wavelength: 842 nm) and B04 (resolution: 10 m/px; central wavelength: 665 nm), respectively.

The metric ΔNDVI was then derived as:

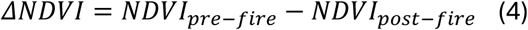

Given that the 2018 fire occurred in late September, NDVI was calculated in early autumns for 2017 to represent pre-fire baseline conditions and for 2018–2024 to assess post-fire vegetation regrowth. Vegetation changes were quantified using ΔNDVI, representing relative gains or losses over time, with higher values indicating more advanced recovery (Bousquet et al., 2022; Lasaponara et al., 2022). NDVI is a bounded index ranging from −1 to +1 and is therefore not expected to follow a Gaussian distribution (Balata et al., 2022). Consequently, NDVI values were extracted using stratified random sampling proportional to land-cover class size (pixel count), with an equal number of pixels randomly sampled from each class to ensure balanced comparisons.

Annual NDVI distributions were generated using a nonparametric bootstrap with 1,000 iterations (Efron et al., 1993) and summarized using box-and-whisker plots. For inter- and intra-class as well as inter-annual comparisons, a single bootstrap realization was selected as a representative NDVI extraction, preserving distributional properties while ensuring consistency across classes and years. These distributions were then summarized for each class for 2017–2024, enabling comparison of post-fire recovery dynamics across vegetation types in burned and unburned areas.

To evaluate if the two distributions come from the same statistical population the Mann–Whitney U test was applied. Differences between burned and unburned areas were considered significant when more than 95% of bootstrap replicates yielded p-values < 0.05.

## 3 Results

### 3.1 LC Model Approaches and Stability Metrics

Over several years, MULTI maintained a categorization accuracy between 70% and 82%. In contrast, the SINGLE approach observed declining overall accuracy in years far from the training one spanning from 63% to 81% (see all results in SM.2 Table S3).

The coefficient of variation (CV) and pixel stability metrics together indicate a high degree of temporal stability across the mapped area. As shown in Table 1, CV patterns differ among classes: grassland, scree, and shrubland exhibit lower CV values under the SINGLE approach than under the MULTI model, whereas woodland classes (T1, T2, T3), olive groves, and needle-leaved shrub (N) consistently show lower CV values under the MULTI approach compared with the SINGLE model.

**Table 1.**
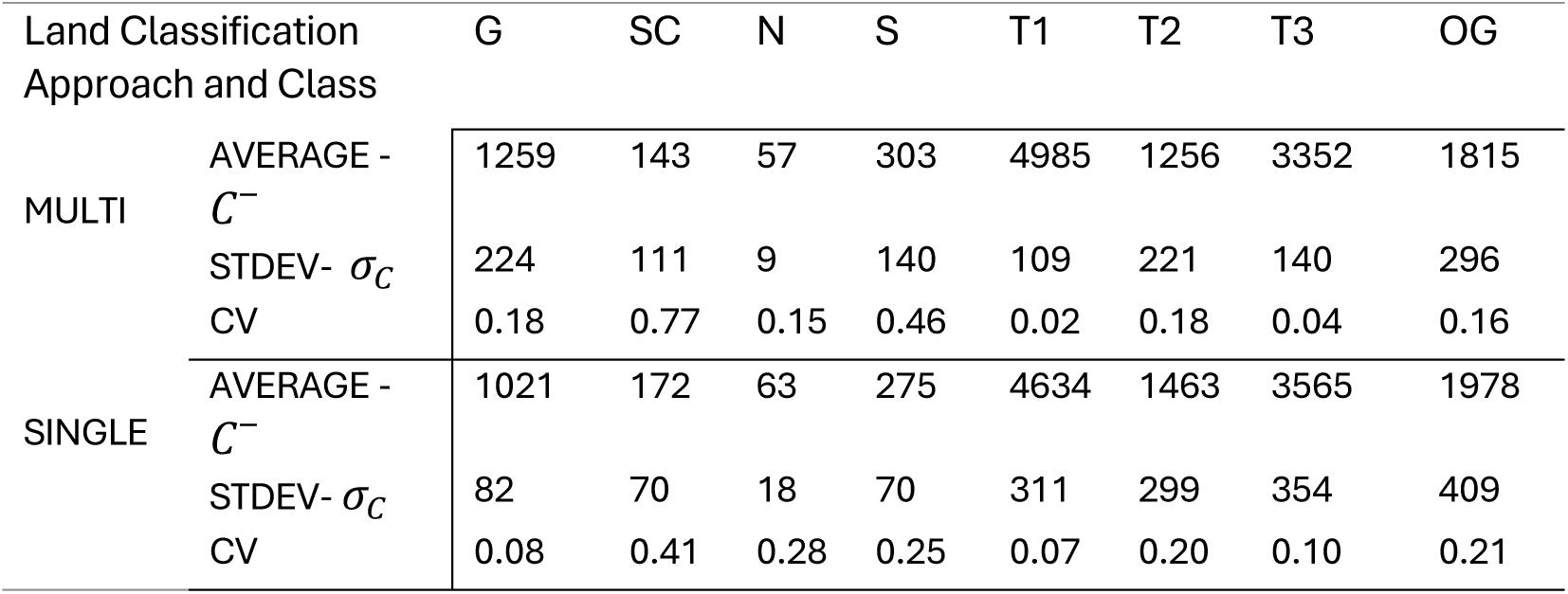
Results of model stability metrics for the study areas, including yearly mean values and standard deviations (in hectares) for all mapped classes under both approaches, together with the corresponding coefficients of variation (CV). Land-cover class acronyms are as follows: (G) grassland, (SC) scree, (N) needle-leaved shrub, (S) shrubland, (T1) broadleaved deciduous forest, (T2) broadleaved evergreen forest, (T3) coniferous forest, and (OG) olive groves.

The results of pixel stability for the MULTI and SINGLE approaches are shown in Figure 2. A clear distinction is observed, with the MULTI models showing a significantly larger unchanged area compared to the SINGLE approach. The MULTI method shows a number of pixels with no change in an eight-years period that is more than the SINGLE approach.

**Figure 2.**
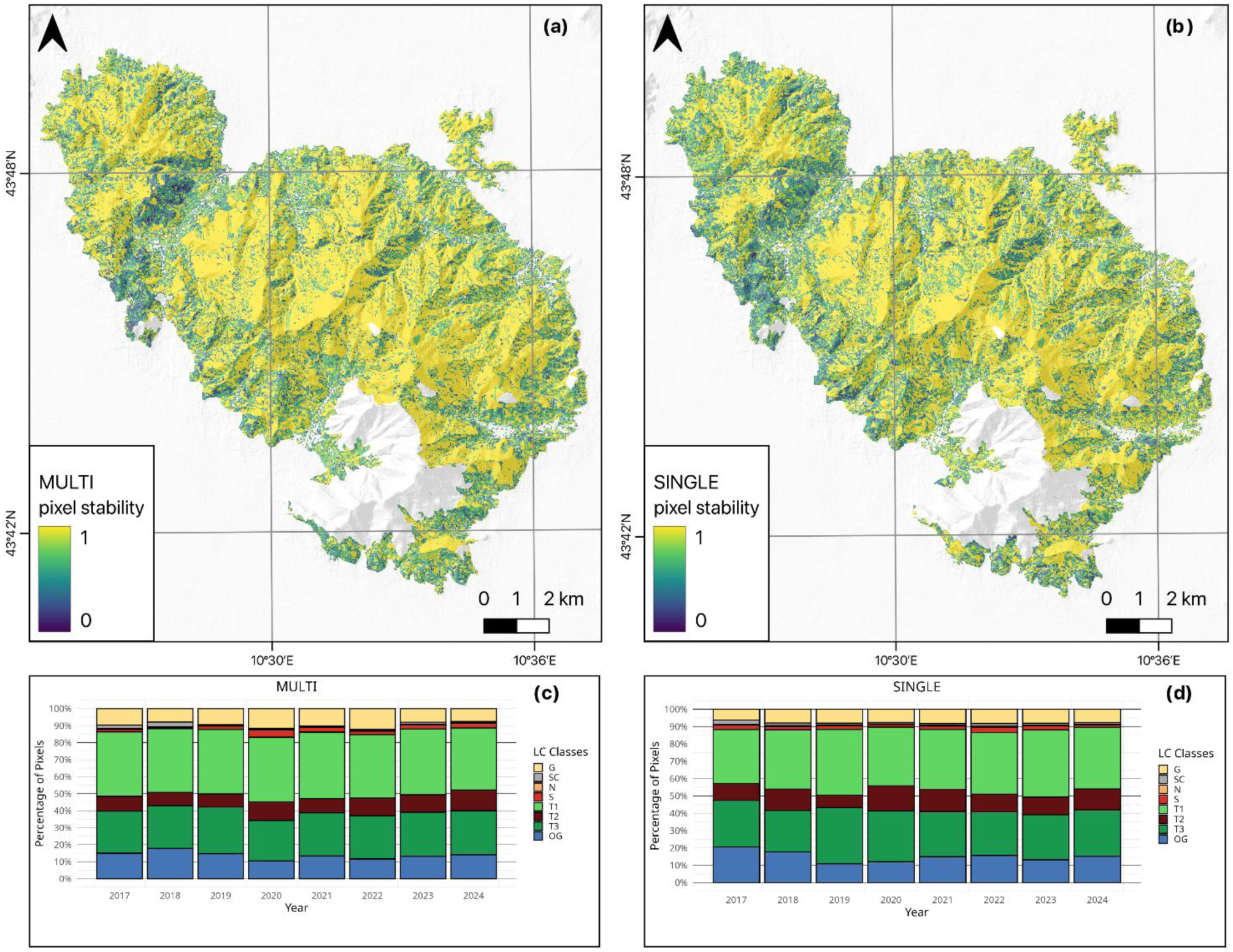
Pixel stability for the MULTI (a) and SINGLE (b) classification approaches together with the pixel variation over time are shown for both the MULTI (c) and SINGLE (d) in the Monte Pisano outside the 2018 Calci fire area (white patch). A value of 1 indicates no temporal change, whereas a value of 0 indicates the maximum temporal change. Land-cover class acronyms are as follows: (G) grassland, (SC) scree, (N) needle-leaved shrub, (S) shrubland, (T1) broadleaved deciduous forest, (T2) broadleaved evergreen forest, (T3) coniferous forest and (OG) olive groves. Class N is not visible due to its small size.

Pixel stability analysis shows a better performance of the MULTI model method compared with the SINGLE approach. Furthermore, the spatial distribution of stability values (panel a) reveals that the most stable pixels (values near 1.00) are more clearly expressed in the MULTI results. These highly stable areas are mainly concentrated in the upper elevations of the Monte Pisano (approximately 400–800 m a.s.l.) and are generally located away from urban and agricultural interface zones.

### 3.2 Post-fire LC classification accuracy

Classification accuracy was assessed through field validation in the Calci fire area using standard accuracy metrics. The model achieved an overall accuracy of 78% (see SM.2 Table S4). Grasslands exhibited the greatest classification uncertainty, primarily due to frequent confusion with olive groves. F1-scores exceeded 0.94 for all classes (macro-F1 = 0.97), and the model achieved an AUC of 0.91, indicating strong predictive performance despite grassland–olive grove overlap. Overall, these results support the robustness of the MULTI approach for mapping land-cover change and post-fire recovery dynamics using remote-sensing data.

### 3.3 Assessment post-fire LC trajectories

Following the comparative evaluation of land-cover classification methods, the MULTI approach was identified as the most suitable for analyzing post-fire LC transitions and ecological trajectories within the burned area. Accordingly, this section presents the results of LC classification and remote-sensing analyses conducted over the monitoring period (2017–2024).

Figure 3 illustrates vegetation classes predicted by the MULTI model within the 2018 fire perimeter over the period 2017–2024. Prior to the fire event, the landscape was dominated by coniferous forests in the central area, with deciduous forests prevailing at higher elevations. Shrubland ecosystems were primarily distributed in the southern sector and along the rural–urban interface (see SM.3 Figure S1).

**Figure 3.**
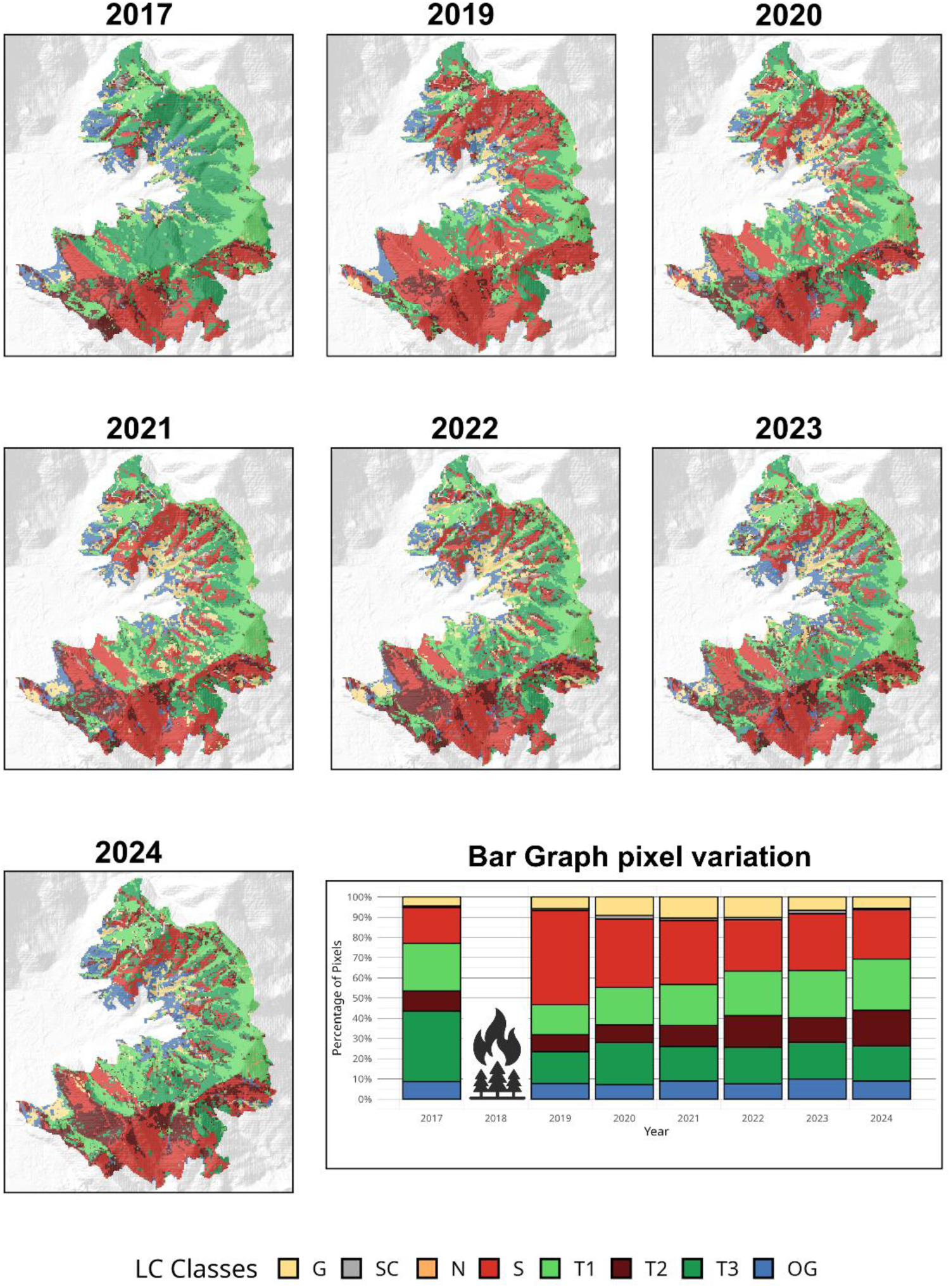
Classification obtained using the MULTI-model approach in the burned area of Monte Pisano–Calci for 2017 (pre-fire) and 2019–2024 (post-fire). The bar chart shows pixel variation (%) over time for land-cover classes (2017–2024). Land-cover class acronyms are as follows: (G) grassland, (SC) scree, (N) needle-leaved shrub, (S) shrubland, (T1) broadleaved deciduous forest, (T2) broadleaved evergreen forest, (T3) coniferous forest, and (OG) olive groves.

After the fire event, the model predominantly classified the burned area as shrubland. In following years, shrubland extent gradually declined, while other vegetation types expanded (see SM.2 Table S5, for total classified pixel counts; n = 28,925 and see SM.2 Table S6, for land-cover transition percentages). Specifically, as shown in Figure 3b and Figure 4, shrub cover increased markedly (>160%) in 2019 compared to pre-fire conditions in 2017. In contrast, deciduous and evergreen broadleaf forests declined by 16–35%, while coniferous cover (primarily *Pinus spp.*) exhibited a more pronounced reduction of approximately 55%.

**Figure 4.**
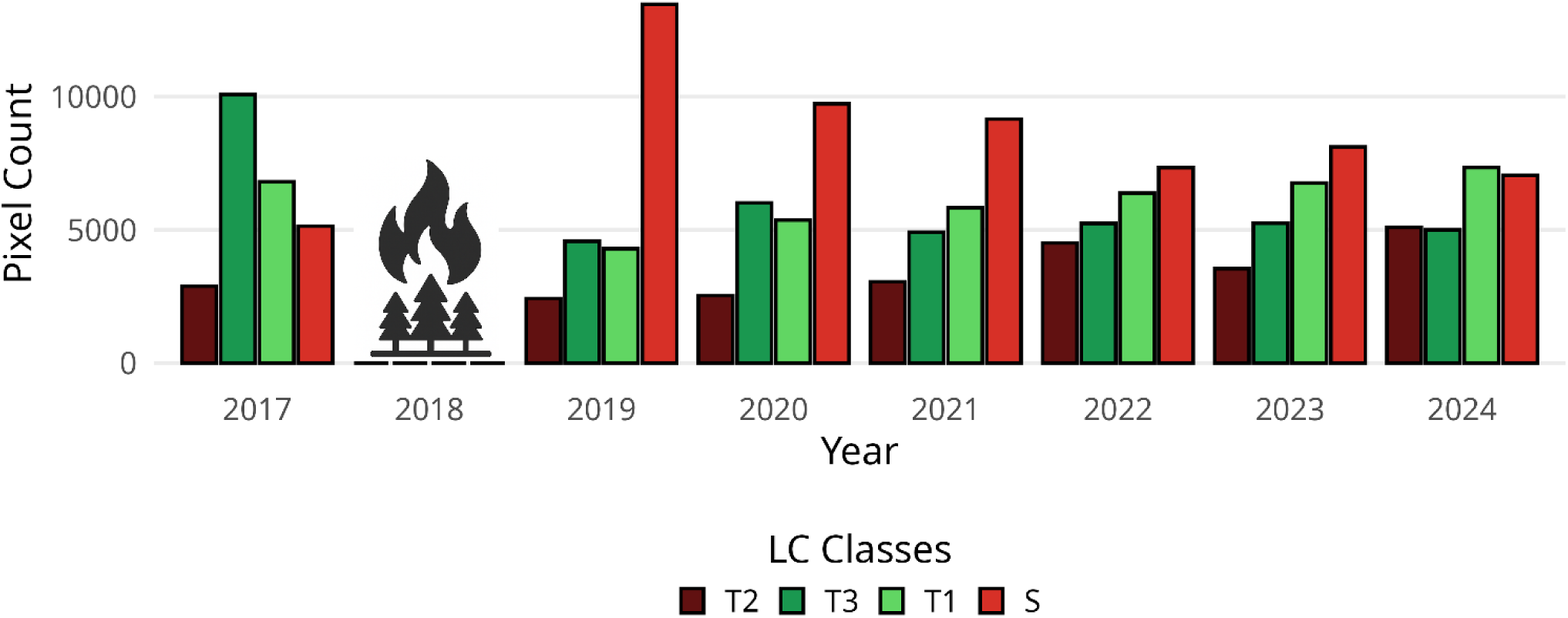
Bar charts showing yearly pixel count variation for forest and shrubland stands from 2017 to 2024, including the 2018 wildfire event. Land-cover class acronyms are as follows: (T1) broadleaved deciduous forest, (T2) broadleaved evergreen forest, (T3) coniferous forest and (S) shrubland.

Furthermore, these vegetation cover trajectories and associated pixel change trends are consistent with the ΔNDVI (Figure 5). Over the examined 2018-2024 period for ΔNDVI, results show a progressive increase in NDVI values during the post-fire period, indicating gradual vegetation recovery within the burned area.

**Figure 5.**
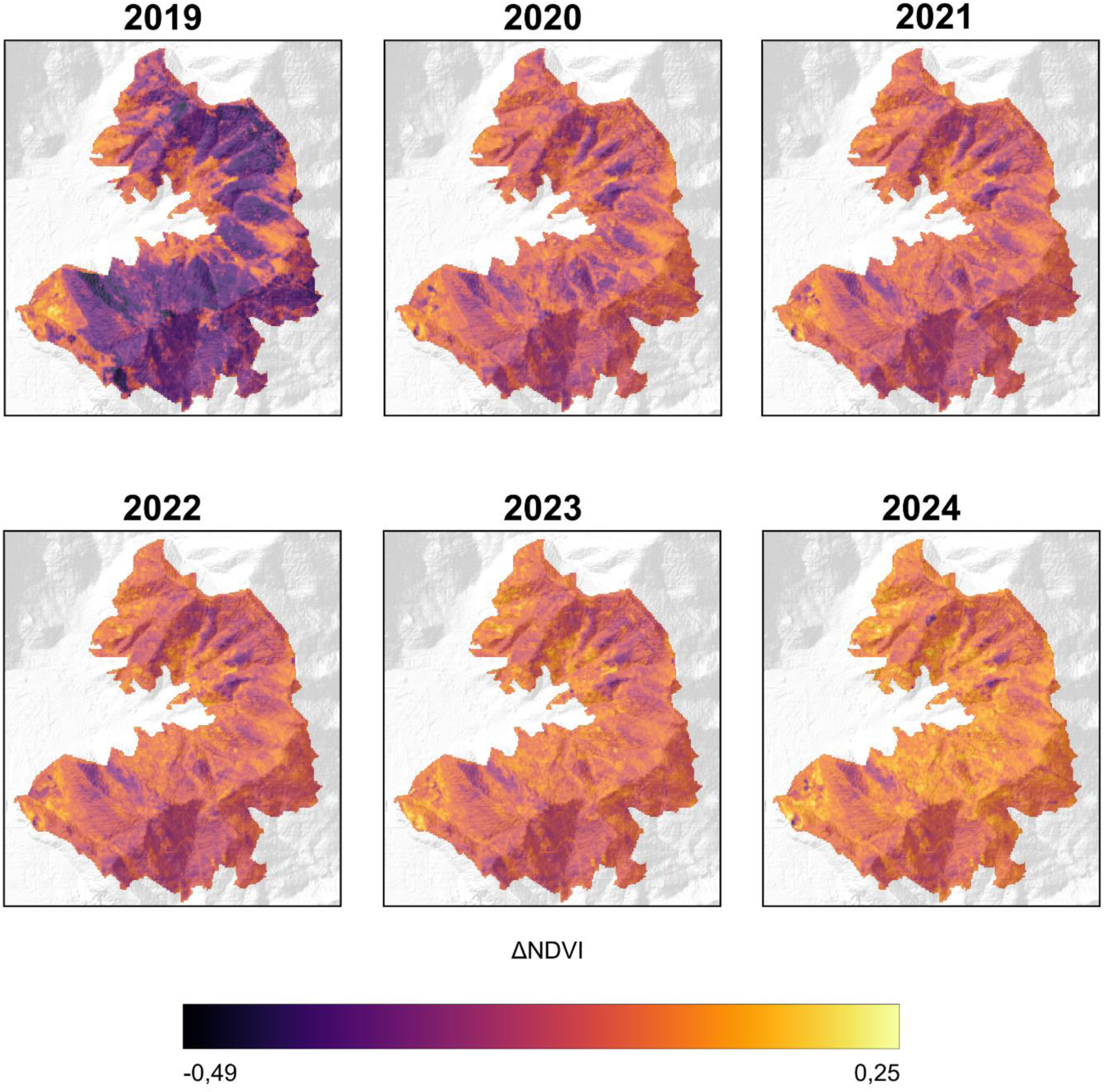
Spatial variation in the 2018 Calci fire area of post-fire vegetation recovery derived from a ΔNDVI time series spanning 2019 to 2024.

Burned areas exhibit pronounced interannual variability in NDVI (Figure 6), consistent with post-fire vegetation regrowth dynamics, whereas unburned areas remain comparatively stable over the study period, showing only minor fluctuations.

**Figure 6.**
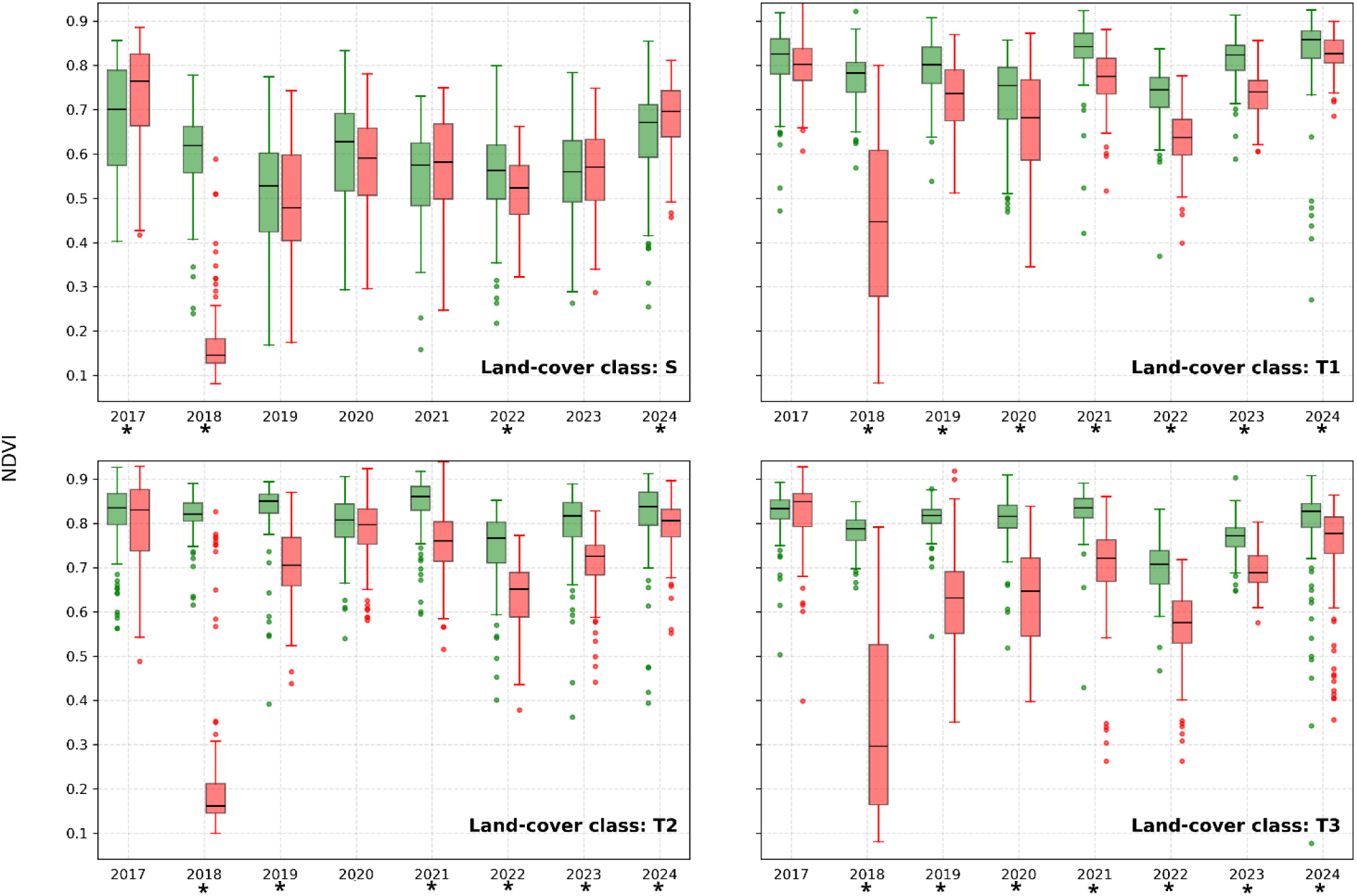
Box-Whisker plots of NDVI distributions from 2017 to 2024 in the unburned (green) and burned (red) areas of the Monte Pisano study site; The symbol * below the years in the x axis denotes distribution difference statistical significance based on the Mann–Whitney U test (p < 0.05). Land-cover class acronyms are as follows: (T1) broadleaved deciduous forest, (T2) broadleaved evergreen forest, (T3) coniferous forest and (S) shrubland.

## 4 Discussion

Previous studies have examined post-fire vegetation regeneration in relation to plant growth strategies and fire severity (Pausas et al., 2014; Zheng et al., 2022). Characterizing post-fire recovery is challenging due to the multiple environmental drivers shaping vegetation regrowth and producing diverse recovery trajectories (Xofis et al., 2021; Blanco-Rodríguez et al., 2023) that can vary across different spatial scales (e.g., Ireland et al., 2015; Viana-Soto, 2017; Loudermilk et al., 2022).

Despite data limitations and the focus on one model study area, our results confirm that satellite-based monitoring and spectral vegetation indices can effectively capture post-fire regrowth, improving classification performance and long-term mapping of vegetation recovery in Mediterranean ecosystems.

### 4.1 Comparing Random Forest classification performance using multi-year versus single-year training models

The performance of the MULTI approach, particularly for tree-cover classes, underscores the importance of explicitly representing interannual variability in vegetation-cover classification for Mediterranean sites. By integrating multi-year information, the model captures climate and phenology-driven shifts in vegetation reflectance (Dronova & Taddeo, 2022) as well as disturbance-induced changes in canopy structure, resulting in more temporally stable classification results. In contrast, uncertainty in the SINGLE approach (i.e. tree-cover classes) increases with temporal distance from the training year, revealing limited temporal transferability. This pattern indicates that even short temporal lags between training and application can introduce bias in model-based estimators (Filippelli, et al 2024). The reduced CV observed in forested ecosystems suggests that the MULTI model more effectively captures ecological stability over time than the SINGLE approach. In contrast, higher variability in shrubland classes likely reflects interannual fluctuations in vegetation growth driven by climatic conditions (Figure 6 see panel LC class S). Our findings demonstrate that incorporating temporal variability is not merely a technical improvement but a prerequisite for reliable land-cover monitoring in disturbed landscapes. Consequently, multi-year training strategies provide a transferable and robust framework for long-term vegetation monitoring and change detection in this Mediterranean site.

### 4.2 Assessing post-fire land-cover trajectories

Although satellite spectral data do not directly represent vegetation regrowth (Pérez-Cabello et al., 2021) - for example, they cannot quantify active above-below ground biomass - recovery trends in vegetation cover can be reliably inferred by linking spectral change metrics with appropriate reference datasets (e.g., ground-based vegetation observations). In this study, time-series satellite data combined with spatially explicit vegetation cover modelling effectively captured the extent and spatial distribution of post-fire land-cover classes. The resulting trajectories revealed both continuous and discontinuous dynamics, enabling the identification of class-specific recovery patterns driven by differences in regrowth strategies, canopy development, and structural change over time (Figure 3).

Our results, as well as our field observations (i.e., ground truth) of vegetation cover collected five years after the wildfire (2023), confirmed the increase of evergreen shrub and tree cover, consistent with both model predictions and temporal trends in NDVI values (Figure 4). These patterns align with established post-fire dynamics in forest ecosystems, where initial recovery is largely driven by resprouting plants (benefitting from resources and buds stored in specialized organs, such as lignotubers; Paula et al., 2016), followed potentially by seed germination (Lamont & Downes, 2011).

Annual transitions in tree and shrub cover within the burned area (Figure 4) are mirrored by corresponding shifts in NDVI values across land-cover classes before and after the 2018 wildfire (Figure 6). These patterns reveal distinct recovery trajectories for tree and shrub communities, possibly driven either by local environmental factors such as soil nutrients and drainage, or by local scale plant-environment (Rietkerk et al 2004) and/or plant-fire feedbacks, that can vary drastically with plant functional type depending e.g. on their post-fire responses (e.g., Baudena et al 2020), rather than by broad-scale environmental drivers.

Areas experiencing minor fire damage (Figure 5) were located at the margins of the Calci wildfire, including interfaces with croplands (e.g., olive groves) and urban zones at the lower elevations, as well as mixed-mesophytic forests dominated by *Castanea sativa* (sweet chestnut) and *Quercus cerris* (Turkey oak) at higher elevations of Monte Pisano. These forest stands are typically found on north-facing slopes with moist, deep soils around 500 m a.s.l., often within runoff-collecting depressions (*impluvium* terraces), which may have contributed to limit the spread and mitigate the intensity of fire.

Satellite-derived ΔNDVI signals indicate that reductions in vegetative vigor largely coincide with areas of highest fire severity, particularly within the 300–350 m a.s.l. belt and on shallow, gravel substrates or exposed bedrock, where pre-fire tree and shrub communities are now beginning to resprout and recolonize (see SM.3, Figure S2). Despite widespread vegetation recovery, areas severely affected by the 2018 fire event remain pronounced in 2024, particularly in the eastern and western sectors formerly dominated by pine forests. These sites experienced extensive canopy loss and currently support only sparse juvenile *Pinus* recruitment (see SM.3, Figure S3). Vegetation is now characterized by transitional shrub assemblages, with co-occurring evergreen broad-leaved (often resprouter) species such as strawberry tree (*Arbutus unedo*), alongside recovering stands of maritime pine (*Pinus pinaster*).

NDVI trends for woodlands and shrublands reveal a sharp decline in tree and shrub vigor in 2018, which—consistent with findings from similar studies in the EU and the United States—represents a clear spectral signal (i.e., a breakpoint) in multitemporal NDVI time series (Gouveia et al., 2010; Bright et al., 2019). Recovery patterns vary with vegetation characteristics associated with plant functional types (Smith et al., 1993), shrublands and broadleaf forests have largely regained pre-fire vigor, whereas coniferous stands exhibit the most persistent decline. As NDVI spectral index does not provide accurate biomass quantification (Fern et al., 2018), the magnitude of losses and the rates of subsequent regrowth for woody and shrubby vegetation affected by the 2018 wildfire remain uncertain.

Furthermore, the NDVI analysis also reveals several “negative outliers” scattered all across the examined time period (Figure 6), indicative of declines in vegetation vigor (“browning”). In unburned areas, these likely reflect plant senescence, pest outbreaks (e.g., insects), silvicultural interventions or drought effects (De Jong et al., 2012). Indeed, severe meteorological drought conditions affecting Europe in 2022, characterized by record-low precipitation and high temperatures (Bevacqua et.al, 2024) likely drove the downward NDVI trajectory between 2021 and 2022 in the study area. A similar pattern is observed in burned areas, especially for tree’s cover (T1-3) from 2021, suggesting the re-establishment of natural processes through vegetation turnover, as observed in undisturbed areas of Monte Pisano (and from field-observations of dead tree remains). Conversely, in 2018, the NDVI for burned area shows “positive outliers” with extended whiskers relative to higher median values across all cover types, reflecting a partial “greening” effect. This pattern suggests rapid regrowth of constant species, likely boosted by post-fire increases in leaf biomass and photosynthetic activity (Kim et al., 2024).

### 4.3 Quantifying vegetation recovery patterns to guide targeted strategies for post-fire Mediterranean landscape resilience and restoration

For 2019–2024, we observed an increase in the area occupied by evergreen tree species (T2), primarily resprouters (Figure 4 and SM.2, Table S6). In contrast, following the post-fire decline, coniferous stands (T3), which are obligate seeders, remained largely stable. Deciduous tree species (T1) exhibited a consistent increase, whereas shrubland (S), hosting several resprouter species, showed a steady decrease. Therefore, the recovery of indigenous tree cover types was observed over the last three years (2022–2024), specifically in classes T1 and T2 (SM.3 Figure S4), corresponding to habitats listed under Directive 92/43/EEC (i.e., 91M0-Pannonian-Balkanic turkey oak-sessile oak forest, 9260-*Castanea sativa* woods and 9340-*Quercus ilex* and *Quercus rotundifolia* forests).These findings provide strong evidence of natural habitat resilience (based on local species-pool filtering) and have important implications for the long-term preservation of the historical landscape of the Monte Pisano, particularly in contrast to the extensive anthropogenic tree (primarily of *Pinus pinaster*) plantations established in the study area over the last ∼80 years (Bertacchi et al., 2004). This is also in line with long-term modelling studies showing that evergreen mesic Mediterranean forests are very resilient to wildfires (Vasques et al 2023), even though they might be destabilized in the future by exacerbated aridity driven by climate change (Baudena et al 2020).

Our results reveal a relationship between fire effects and ecosystem regeneration, with vegetation-cover transitions driven by plant regrowth mechanisms that promote rapid post-fire recovery. Findings also suggest that environmental factors (e.g., solar radiation and soil moisture) and ecological processes (e.g., plant competition, species pool filtering; Ladouceur et al., 2018), and plant functional type composition (Kattenborn et al., 2019) can contribute to shaping vegetation recovery rates (Ireland et al., 2015; Viana-Soto, 2017; Loudermilk et al., 2022). In Mediterranean landscapes, north-facing and gentle slopes often exhibit higher regeneration rates following disturbance due to reduced evapotranspiration, greater moisture availability, and enhanced soil-nutrient retention (Röder et al., 2008). Additionally, resprouting species generally support more rapid recovery than obligate seeders (Pausas et al., 2014).

In our study area, post-fire vegetation dynamics is undergoing natural recolonization, with successional trajectories potentially directing toward forest and shrubland ecosystems consistent with the Habitat Directive (92/43/EEC). This process should be supported not only through national and regional management interventions, but also by recognizing the inherent ecological resilience of Mediterranean species, which have shaped both southern and northern Mediterranean landscapes since the Miocene. Such recognition can inform restoration strategies that identify and prioritize species resilient to current and predicted climatic conditions.

## 5 Conclusions

In conclusion, the proposed machine learning–based framework enables spatially explicit monitoring of post-fire vegetation dynamics and robust comparison of recovery trajectories across Mediterranean fire-prone ecosystems. The integration of remote-sensing spectral index time series strengthens landscape-scale assessments of post-fire recovery, while future incorporation of fine-scale environmental drivers may further enhance machine learning models performance.

The framework supports targeted restoration actions aligned with the European Nature Restoration Regulation (EU-2024/1991) and contributes to United Nations Sustainable Development Goal 15 (Life on Land) by: (i) prioritizing Mediterranean species adapted to current climatic conditions; (ii) supporting binding post-fire restoration strategies that favor natural ecosystem recovery over afforestation with non-native species; (iii) enhancing biodiversity conservation at both species and genetic levels; and (iv) promoting the long-term climate resilience of Mediterranean woodlands.

Overall, the results demonstrate that reinforcing recovery processes governed by native vegetation dynamics can enhance ecosystem resilience, functional integrity, and long-term climate stability.

## Glossary

The following abbreviations are used in this manuscript:

CV: Coefficient of Variation
ÄNDVI: delta-Normalized Difference Vegetation Index
LC: Land-Cover
NDVI: Normalized Difference Vegetation Index
RF: Random Forest
S2: Sentinel 2 MSI

## CRediT authorship contribution statement

Emiliano Agrillo: Writing – review & editing, Writing – original draft, Validation, Resources, Methodology, Conceptualization. *Equal Contribution*.

Nazario Tartaglione: Writing – review & editing, Writing – original draft, Validation, Software, Resources, Methodology, Data curation, Conceptualization. *Equal Contribution*.

Alessandro Mercatini: Writing – review & editing, Writing – original draft, Software, Methodology, Data Curation, Resources.

Alice Pezzarossa: Writing – review & editing, Writing – original draft, Validation, Methodology. Gianluigi Ottaviani: Writing – review & editing

Mara Baudena: Writing – review & editing

Federico Filipponi: Writing – review & editing, Resources, Methodology, Data curation.

## Acknowledgements

The authors affiliated with ISPRA (EA, NT, AM and AP) gratefully acknowledge the support provided by the organization in carrying out this research. Other authors (GO, MB and FF) acknowledge the Italian National Biodiversity Future Center (NBFC): National Recovery and Resilience Plan (NRRP), Mission 4 Component 2 Investment 1.4 of the Italian Ministry of University and Research; funded by the European Union - NextGenerationEU (Project code CN_00000033).

## Supporting Information

Additional supporting material can be found online in the Supporting Information section at the end of this article: i) SM.1 - Description of features used in the Random Forest models, Table 1 List of predictor variables (Environmental & Spectral) used for land-cover classification modelling, including corresponding metadata descriptions, Table 2 List spectral indices adopted, equation based on Sentinel-2 Multispectral Instrument bands, and references; ii) SM.2 - Table 3 Overall accuracy (OA) and its standard error (SE) for the MULTI and SINGLE approach computed on the predicted classes, Table 4 Contingency table comparing predicted and observed land-cover classes within the wildfire area (year 2023), Table 5 Predicted land-cover classes within the burned area, expressed as pixel counts (total area = 28,925 pixels), Table 6 Land-cover transition pixels classified (in %), within the wildfire area, compared to 2017 (pre-fire); iii) SM.3 – Photographs *FigureS1, FigureS2, FigureS3, FigureS4*.

## Notes

### Competing Interest Statement

The authors have declared no competing interest.

