## Supplementary material for "ASSESSING POST-FIRE VEGETATION TRAJECTORIES USING MACHINE LEARNING AND REMOTE SENSING: EVIDENCE FROM A MEDITERRANEAN SITE": Supp

- **SM.1: Description of features used in the Random Forest models (lists, formulas, and sources), Tables S1 & S2**
- **SM.2: Tables 3-6**
- **SM.3: Photographs, Figures 1-4**

***

- SM.1: Description of features used in the Random Forest models

The predictors are grouped into two main categories:

1. Environmental data: variables mainly related to geographic, topographic, climatic, and soil characteristics.
2. Spectral data: values acquired from the Multi Spectral Instrument (MSI) onboard Sentinel-2 satellites.

All variables were obtained or derived from open-access datasets and resampled to a consistent 20 m spatial resolution (see SM.1 Table 1 list of all predictor variables).

The reference spatial grid, with coordinate reference system EPSG:32632, aligns with the matches the coordinate system of the T32TPP tile covering our ROI Sentinel-2 MSI satellite imagery available over the study area. We chose a grid with 20 m grid spacing because it represents the plurality of Sentinel-2 spectral surface reflectance bands resolutions (four bands are 10 m, six are 20 m, and three are 60 m). Resampling all bands to 20 m ensures consistency and compatibility across the entire dataset.

Environmental Variables

Environmental variables were selected based on their relevance to plant dispersal processes. These variables are assumed to be time-invariant (Immitzeret al., 2019; Descombes et al., 2020). Among geographic variables, distances from both the seashore and the river network were calculated for each grid cell, emphasizing shoreline proximity. Topographic features, like slope and aspect, were derived from a 20 m resolution Digital Elevation Model (DEM) of Italy, using a 3 × 3 pixel moving window. From the aspect values, the northernness and easternness components were computed using sine and cosine transformations to generate continuous variables. Climatic variables included time average precipitation and temperature obtained by ground stations and averaged over time at 1 km horizontal resolution (Fioravanti et al., 2010; Braca et al., 2018). These were interpolated to the 20 m reference grid using a regularized spline with tension. To standardize temperature values and eliminate elevation-related bias, an altitude correction was applied (US standard atmosphere NOA, 1976). The daily average solar irradiance was calculated at a spatial resolution of 20 m using equations for solar energy-related characteristics, taking local topography into account (Súri and Hofierka, 2004). Then it was compiled to determine monthly averages, while daytime cloud cover obtained from the ESA–Climate Change Initiative cloud dataset for the period 2004–2014 was utilized to compute the monthly average climatological cloud cover percentage (Duvellier et al. 2021). These averages were resampled using a regularized spline to match the 20 m resolution keeping into account an estimation of sky diffuse radiation (Karsten et al., 1980). The result was a monthly weighted solar irradiance dataset used to compute daily averages. Soil properties, specifically pH and depth to bedrock, were extracted from the SoilGrids dataset, originally at 250 m resolution, and interpolated to the reference grid using a Bartlett filter with a 750 m radius for spatial consistency.

Spectral Variables

Spectral data included surface reflectance bands and derived spectral indices from Sentinel–2 MSI. This sensor offers high spatial resolution, a five-day revisit time, a 290 km swath width, and a spectral range spanning from visible to shortwave infrared. All Sentinel–2 images acquired from 2017 to 2024 with cloud cover below 90% were collected. While this threshold may seem high, it is essential for obtaining reliable composite images based on monthly averages. Each valid observation within a given tile contributed to the analysis, whereas cloud-covered areas bad pixels (namely pixels flagged as cloud, cloud shadow, snow, topographic shadow) were excluded from the calculation of monthly averages. The use of multitemporal data time series in land cover classification is more effective than relying on single-date results. The surface reflectance bands were adjusted to a size of 20 m, resulting in a consistent dataset that includes the following bands: B2 (blue), B3 (green), B4 (red), B5, B6, B7, B8A (Red Edge), B8 (near infrared 1), B11 (short-wavelength infrared 1), and B12 (short-wavelength infrared 2). Any missing data at the pixel level caused by persistent cloud cover within a month was imputed using the *imputeTS* R package (Moritz and Bartz-Beielstein, 2017). Along with the Sentinel–2 MSI spectral bands, four other spectral indices (EVI, NDYI, RI, and CRI1 as described in SM.1 Table 2 below) were selected to examine the green, red, and yellow pigments in leaves during blooming and senescence, which is the time when the plant stops making chlorophyll, which shows different accessory pigments. The final classification was based on the complete set of monthly-frequency annual time series data from the Sentinel–2 MSI bands and the four spectral indices like in Agrillo et al. (2021).

*SM.1 Table S1 - List of predictor variables (Environmental & Spectral) used for land-cover classification modelling, including corresponding metadata descriptions.*

| GROUP | SUBGROUP | FIELD_NAME | UNITS | DESCRIPTION | SOURCE |
|---|---|---|---|---|---|
| **Environmental predictors** | *Geomorphologic* | elevation | m a.s.l. | Elevation | ISPRA Database (https://www.isprambiente.gov.it/it/banche-dati) |
| | | slope | degrees | Slope | ISPRA_CSA_data repository |
| | | eastness | Polar units | Eastness | ISPRA_CSA_data repository |
| | | northness | Polar units | Northness | ISPRA_CSA_data repository |
| | *Climatic* | TannNorm | Celsius degrees | Normalized annual average air temperature on elevation | ISPRA Database (https://scia.isprambiente.it/dati-e-indicatori/) |
| | | CRF | mm/year | Annual Cumulated Rainfall | ISPRA Database (https://scia.isprambiente.it/dati-e-indicatori/) |
| | | SolRad | $WH/m^2$ | Daily average solar radiation | ISPRA Database (https://www.isprambiente.gov.it/it/banche-dati) |
| | | SCD | days | Snow Cover Duration | https://www.theia-land.fr/en/product/snow/ |
| | *Geographic* | dCoastLog | log10(m) | Distance from shoreline (Log10) | https://gn.mase.gov.it/portale/servizio-di-consultazione-wms |
| | | dRivLog | log10(m) | Distance from inland waters (Log10) | https://gn.mase.gov.it/portale/servizio-di-consultazione-wms |
| | | CCLAT | - | of the Cell Centroid | ISPRA_CSA_data repository |
| | *Soil properties* | PHIHOX | pH | pH index measured in water solution | SoilGrids -https://soilgrids.org |

| | | | | | |
|---|---|---|---|---|---|
| | | BDTICM | cm | Absolute depth to bedrock | SoilGrids -https://soilgrids.org |
| | *Vegetation properties* | TCD | percentage | Tree Cover Density | https://land.copernicus.eu/ |
| **Spectral predictors** | *Spectral signature* | S2_20m_B2 | reflectance | Sentinel-2 MSI B2 value at yearly month 01 -12 | ISPRA_CSA_data repository |
| | | S2_20m_B3 | reflectance | Sentinel-2 MSI B3 value at yearly month 01 -12 | ISPRA_CSA_data repository |
| | | S2_20m_B4 | reflectance | Sentinel-2 MSI B4 value at yearly month 01 -12 | ISPRA_CSA_data repository |
| | | S2_20m_B5 | reflectance | Sentinel-2 MSI B5 value at yearly month 01 -12 | ISPRA_CSA_data repository |
| | | S2_20m_B6 | reflectance | Sentinel-2 MSI B6 value at yearly month 01 -12 | ISPRA_CSA_data repository |
| | | S2_20m_B7 | reflectance | Sentinel-2 MSI B7 value at yearly month 01 -12 | ISPRA_CSA_data repository |
| | | S2_20m_B8 | reflectance | Sentinel-2 MSI B8 value at yearly month 01 -12 | ISPRA_CSA_data repository |
| | | S2_20m_B8A | reflectance | Sentinel-2 MSI B8A value at yearly month 02 -11 | ISPRA_CSA_data repository |
| | | S2_20m_B11 | reflectance | Sentinel-2 MSI B11 value at yearly month 01 -12 | ISPRA_CSA_data repository |
| | | S2_20m_B12 | reflectance | Sentinel-2 MSI B12 value at yearly month 01 -12 | ISPRA_CSA_data repository |
| | *Spectral index* | EVI | dimensionless | Sentinel-2 MSI EVI value at yearly month 01 - 12 | ISPRA_CSA_data repository |
| | | NDYI | dimensionless | Sentinel-2 MSI NDYI value at yearly month 01 - 12 | ISPRA_CSA_data repository |
| | | RI | dimensionless | Sentinel-2 MSI RI value at yearly month 01 - 12 | ISPRA_CSA_data repository |
| | | CRI1 | dimensionless | Sentinel-2 MSI CRI1 value at yearly month 01 - 12 | ISPRA_CSA_data repository |

*SM1 Table S2 - List spectral indices adopted, equation based on Sentinel-2 Multispectral Instrument bands, and references.*

| SPECTRAL INDEX | EQUATION | REFERENCE |
|---|---|---|
| Enhanced Vegetation Index - **EVI** | $\frac{2.5 \times (B8 - B4)}{(B8 + 6 \times B4 - 7.5 \times B2 + 1)}$ | (Huete et al., 1994) |
| Normalized Difference Yellow Index - **NDYI** | $\frac{(B3 - B2)}{(B3 + B2)}$ | (Sulik & Long, 2016) |
| Normalized Difference Red/Green Redness Index - **RI** | $\frac{(B4 - B3)}{(B4 + B3)}$ | (Escadafal & Huete, 1991) |
| Carotenoid Reflectance Index - **CRI1** | $\frac{(1 \div B2)}{(1 \div B3)}$ | (Gitelson et al., 2002) |

- <u>SM.2: Tables</u>

*SM.2 Table S3 - Overall accuracy (OA) and its standard error (SE) for the MULTI and SINGLE approach computed on the predicted classes.*

| YEAR | MULTI OA(SE) | SINGLE OA(SE) |
|---|---|---|
| 2017 | 0.70(0.07) | 0.67(0.05) |
| 2018 | 0.71(0.06) | 0.63(0.07) |
| 2019 | 0.78(0.06) | 0.65(0.06) |
| 2020 | 0.74(0.07) | 0.70(0.06) |
| 2021 | 0.79(0.06) | 0.81(0.06) |
| 2022 | 0.82(0.06) | 0.79(0.06) |
| 2023 | 0.79(0.06) | 0.79(0.06) |
| 2024 | 0.75(0.06) | 0.75(0.06) |

*SM.2 Table S4 - Contingency table comparing predicted and observed land-cover classes within the wildfire area (year 2023). Land-cover class acronyms are as follows: grassland (G), scree (SC), needle-leaved shrub (N), shrubland (S), broadleaved deciduous forest (T1), broadleaved evergreen forest (T2), coniferous forest (T3), and olive groves (OG).*

| | | **Observed** | | | | | | | | |
|---|---|---|---|---|---|---|---|---|---|---|
| | **LC Class** | **G** | **SC** | **N** | **S** | **T1** | **T2** | **T3** | **OG** | **Total Observ.** |
| **Predicted** | **G** | **1** | - | - | - | 1 | - | - | - | 2 |
| | **Sc** | - | **2** | - | - | - | - | - | - | 2 |
| | **N** | - | - | **1** | - | - | - | - | - | 1 |
| | **S** | - | - | - | **26** | 1 | 1 | 1 | - | 29 |
| | **T1** | - | - | - | - | **17** | - | 2 | - | 19 |
| | **T2** | - | - | - | 1 | 1 | **1-** | 3 | - | 15 |
| | **T3** | - | - | - | 1 | 3 | 1 | **12** | - | 17 |
| | **OG** | 6 | - | - | - | - | - | - | **1-** | 16 |
| | **Total Predict** | 7 | 2 | 1 | 28 | 23 | 12 | 18 | 10 | **101** |

*Overall Accuracy 78 %*

*SM.2 Table S5 – Predicted land-cover classes within the burned area, expressed as pixel counts (total area = 28,925 pixels). Land-cover class acronyms are as follows: grassland (G), scree (SC), needle-leaved shrub (N), shrubland (S), broadleaved deciduous forest (T1), broadleaved evergreen forest (T2), coniferous forest (T3), and olive groves (OG).*

| LC Class | 2017 | 2018 | 2019 | 2020 | 2021 | 2022 | 2023 | 2024 |
|---|---|---|---|---|---|---|---|---|
| **G** | *1292* | **wildfire** | 1659 | 2636 | 3020 | 2925 | 1904 | 1616 |
| **SC** | *175* | | 266 | 516 | 287 | 297 | 449 | 165 |
| **N** | *38* | | 31 | 45 | 58 | 51 | 46 | 55 |
| **S** | *5138* | | 13463 | 9733 | 9159 | 7335 | 8110 | 7043 |
| **T1** | *6804* | | 4293 | 5368 | 5828 | 6379 | 6759 | 7342 |
| **T2** | *2885* | | 2420 | 2536 | 3054 | 4505 | 3549 | 5095 |
| **T3** | *10081* | | 4571 | 6011 | 4908 | 5239 | 5248 | 4999 |
| **OG** | *2512* | | 2222 | 2080 | 2611 | 2194 | 2860 | 2610 |

*SM.2 Table S6 - Land-cover transition pixels classified (in %), within the wildfire area, compared to 2017 (pre-fire). Land-cover class acronyms are as follows: grassland (G), scree (SC), needle-leaved shrub (N), shrubland (S), broadleaved deciduous forest (T1), broadleaved evergreen forest (T2), coniferous forest (T3), and olive groves (OG).*

| LC CLASS | 2017 PIXEL | 2018 | 2019 % | 2020 % | 2021 % | 2022 % | 2023 % | 2024 % |
|---|---|---|---|---|---|---|---|---|
| **G** | *1292* | **wildfire** | +28 | +104 | +134 | +126 | +47 | +25 |
| **SC** | *175* | | +52 | +195 | +64 | +70 | +157 | -6 |
| **N** | *38* | | -18 | +18 | +53 | +34 | +21 | +45 |
| **S** | *5138* | | +162 | +89 | +78 | +43 | +58 | +37 |
| **T1** | *6804* | | -37 | -21 | -14 | -6 | -1 | +8 |
| **T2** | *2885* | | -16 | -12 | +6 | +56 | +23 | +77 |
| **T3** | *10081* | | -55 | -40 | -51 | -48 | -48 | -50 |
| **OG** | *2512* | | -12 | -17 | +4 | -13 | +14 | +4 |

- <u>SM.3: Photographs</u>

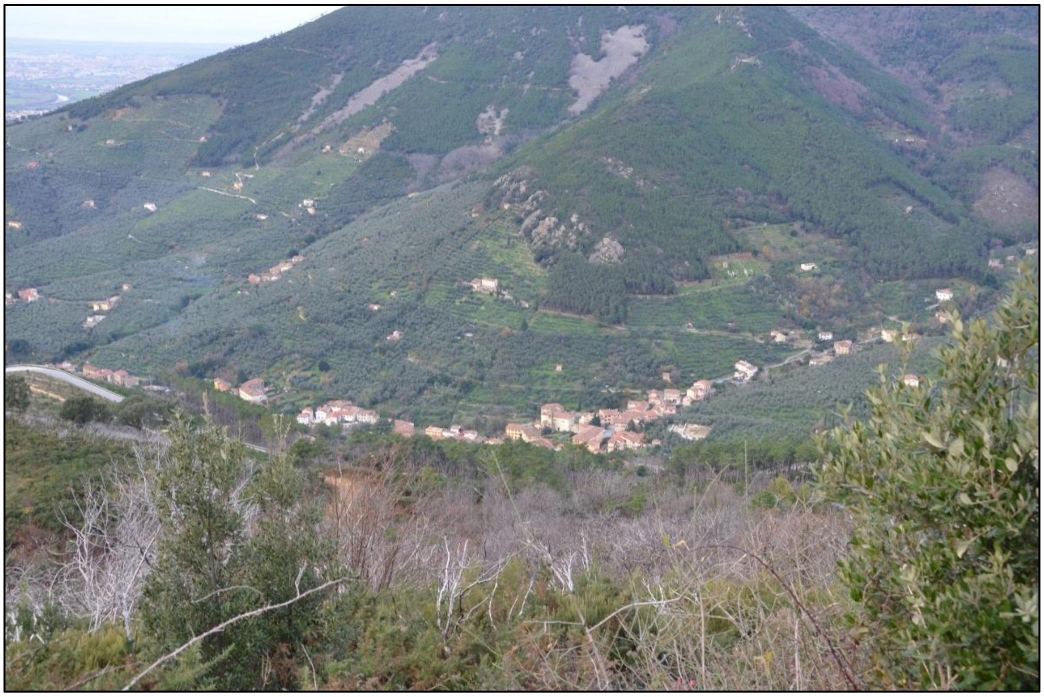

*SM.3 FigureS1 – View of Calci town and the south-facing hillside slope unaffected by wildfires, showing the presence of extensive olive grove patches (photograph taken in 2024 by Nazario Tartaglione).*

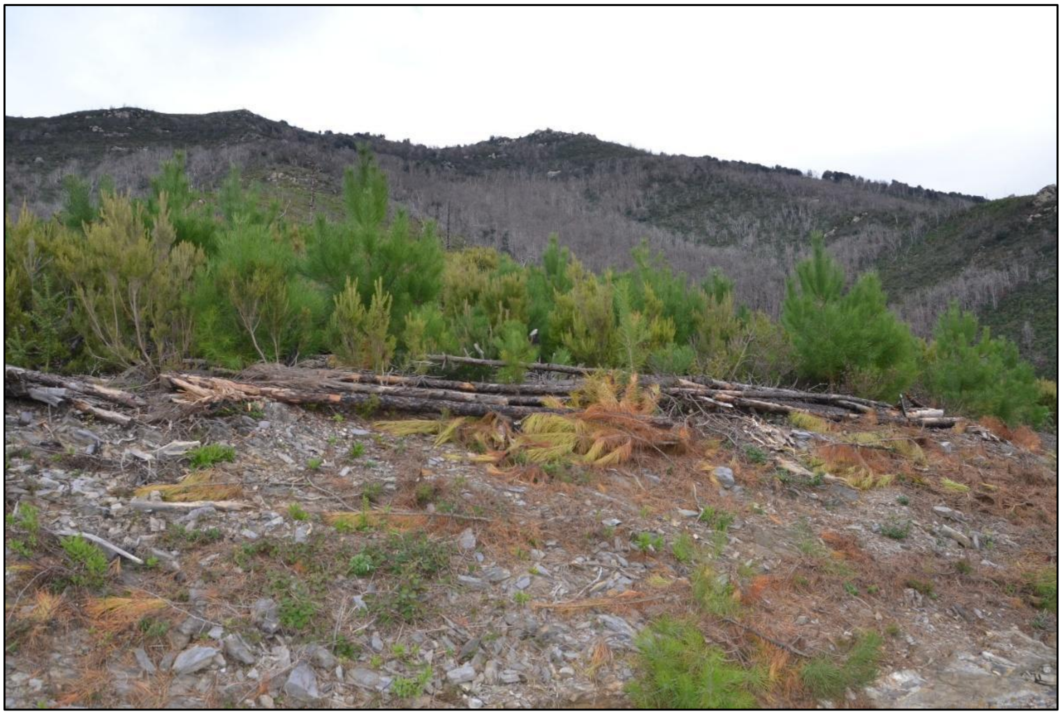

*SM.3 Figure S2 - Post-fire stand showing remains of dead trees affected by wildfire and the establishment of juvenile pine individuals (photograph taken in 2024 by Nazario Tartaglione).*

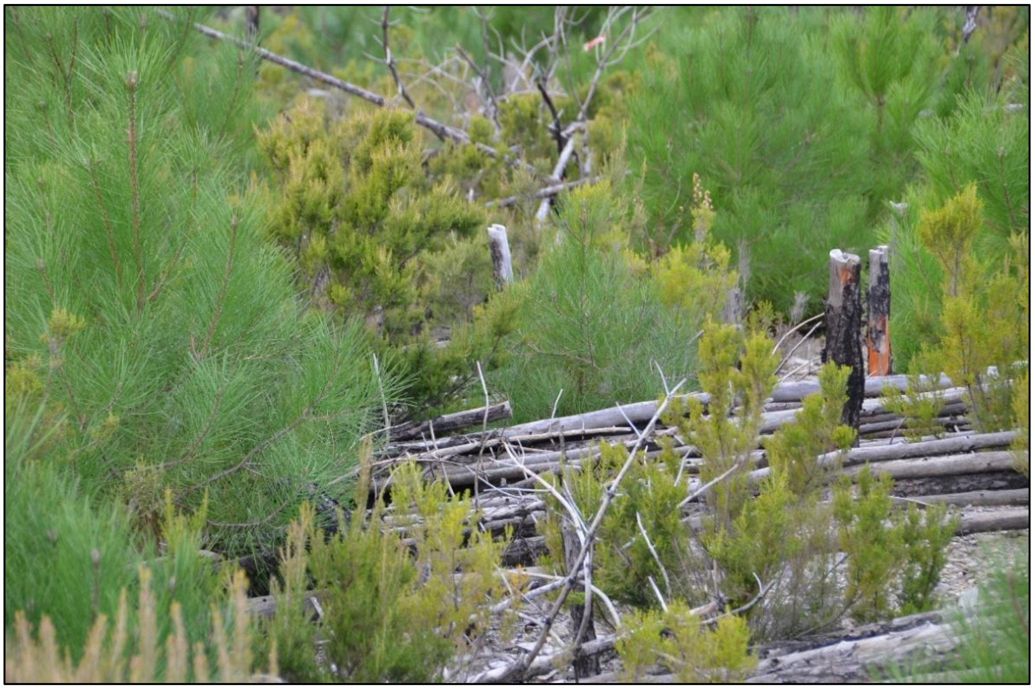

*SM.3 Figure S3 - Juvenile regrowth of pine tree and tree heath observed during post-fire recovery in the Calci area (photograph taken in 2024 by Nazario Tartaglione).*

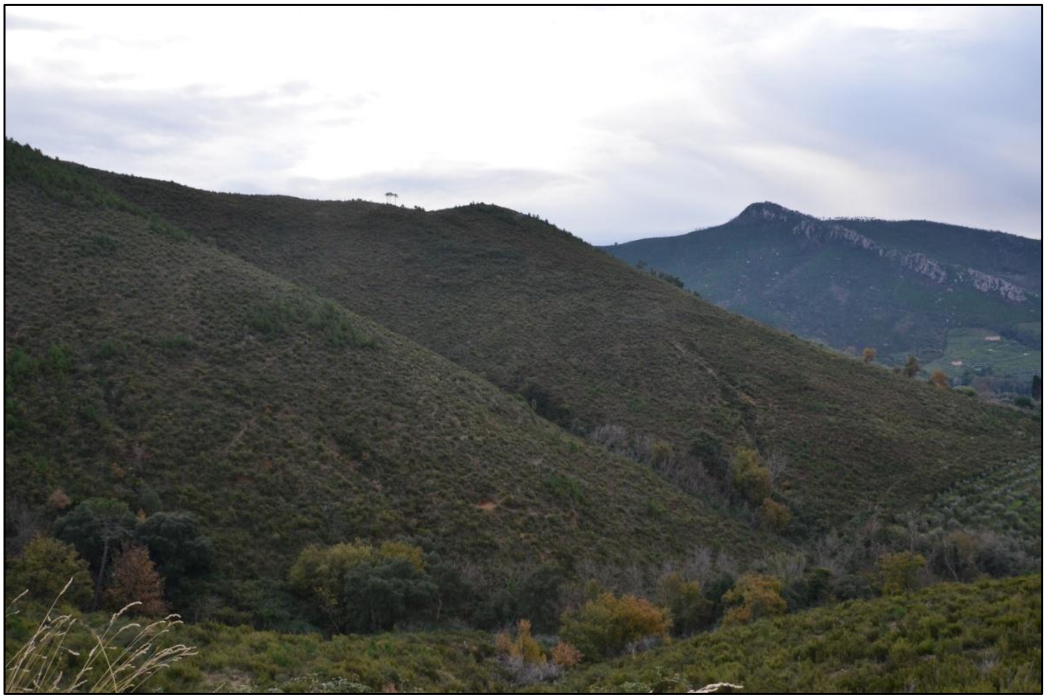

*SM.3 Figure S4 – Post-fire landscape affected by the 2018 wildfire, showing substantial regrowth of evergreen shrublands and juvenile holm oak (Quercus ilex) trees (photograph taken in 2024 by Nazario Tartaglione).*
